# Automated high-throughput biological sex identification from archaeological human dental enamel using targeted proteomics

**DOI:** 10.1101/2024.02.20.581140

**Authors:** Claire Koenig, Patricia Bortel, Ryan S. Paterson, Barbara Rendl, Palesa P. Madupe, Gaudry B. Troché, Nuno Vibe Hermann, Marina Martínez de Pinillos, María Martinón-Torres, Sandra Mularczyk, Marie Louise Schjellerup Jørkov, Christopher Gerner, Fabian Kanz, Ana Martinez-Val, Enrico Cappellini, Jesper V. Olsen

**Affiliations:** Novo Nordisk Foundation Center for Protein Research, Proteomics Program, Faculty of Health and Medical Sciences, University of Copenhagen, Copenhagen, Denmark; Department of Analytical Chemistry, Faculty of Chemistry, University of Vienna, Waehringer Str.38, 1090 Vienna, Austria; Vienna Doctoral School in Chemistry (DoSChem), University of Vienna, Waehringer Str. 42, 1090 Vienna, Austria; Geogenetics Section, Globe Institute, University of Copenhagen, Denmark; Center for Forensic Medicine, Medical University of Vienna, Vienna, Austria; Pediatric Dentistry and Clinical Genetics, Department of Odontology, University of Copenhagen, Copenhagen, Denmark; Centro Nacional de Investigación sobre la Evolución Humana (CENIEH), Paseo Sierra de Atapuerca 3, Burgos, 09002 Spain; Department of Anthropology, University College London (UCL), 14 Taviton Street, London, WC1H 0BW United Kingdom; Laboratory of Biological Anthropology, Globe Institute, University of Copenhagen, Copenhagen, Denmark; Joint Metabolome Facility, University of Vienna and Medical University of Vienna, Waehringer Str.38, 1090 Vienna, Austria; Centro Nacional de Investigaciones Cardiovasculares Carlos III (CNIC), Madrid 28029, Spain

## Abstract

Biological sex is key information for archaeological and forensic studies, which can be determined by proteomics. However, lack of a standardised approach for fast and accurate sex identification currently limits the reach of proteomics applications. Here, we introduce a streamlined mass spectrometry (MS)-based workflow for determination of biological sex using human dental enamel. Our approach builds on a minimally invasive sampling strategy by acid etching, a rapid online liquid chromatography (LC) gradient coupled to high-resolution parallel reaction monitoring assay allowing for a throughput of 200 samples-per-day with high quantitative performance enabling confident identification of both males and females. Additionally, we have developed a streamlined data analysis pipeline and integrated it into an R-Shiny interface for ease-of-use. The method was first developed and optimised using modern teeth and then validated in an independent set of deciduous teeth of known sex. Finally, the assay was successfully applied to archaeological material, enabling the analysis of over 300 individuals. We demonstrate unprecedented performance and scalability, speeding up MS analysis by tenfold compared to conventional proteomics-based sex identification methods. This work paves the way for large-scale archaeological or forensic studies enabling the investigation of entire populations rather than focusing on individual high-profile specimens.

## Introduction

The field of proteomics has gained substantial recognition in archaeological applications, with numerous studies highlighting its potential ^1–5^ by offering a unique window to the past. Using liquid chromatography coupled to tandem mass spectrometry (LC-MS/MS), it is possible to sequence complex peptide mixtures extracted from bio-archaeological material with high sensitivity, enabling the study of extinct organisms ^6^, human evolution ^7^, cultural practices ^3,8^, ancient diets ^9^, population health and disease ^10,11^ or even artistic production techniques ^12–14^.

Characterisation of the biological sex of human remains is a crucial aspect in archaeological and forensics contexts, traditionally relying on the observation of skeletal morphology ^15^ or contextual clues such as the presence of tools or artifacts associated with the human remains ^16^. However, these methods have significant limitations, as they are primarily applicable to adults who have developed clear sexual dimorphic traits ^17^. Similarly, sex inference based on the assessment of contextual material can be affected by historical cultural bias. Molecular evidence was first used for accurate characterisation of the biological sex of ancient specimens through DNA sequencing ^18^, later followed by protein analysis ^19–21^. Applied to archaeological samples, molecular evidence has been able to challenge cultural beliefs about gender and their role and status in past societies ^16,22,23^ as well as interpersonal relations ^24^. However, ancient DNA analysis currently requires at least tens of milligrams of starting bone material, and an elaborate and expensive experimental approach ^25^. Proteomics analysis, although it can be performed micro-destructively using dental enamel acid etching ^20^, currently can only sustainably be applied to small sample sets.

Enamel, the hardest and most mineralised tissue in the human body, containing only 1 to 2% of organic material ^26^, has emerged as an excellent mineral matrix for preserving ancient biomolecules ^27^. Its high degree of mineralisation significantly favours the preservation of proteins ^28^, making it a valuable resource for deep-time phylogenetic studies ^6,7,29,30^. Enamel does not contain DNA, but it contains proteins with a significantly slower degradation rate compared to DNA ^28^. The enamel proteome consists of a low number of highly tissue-specific proteins including amelogenin (AMELX/AMELY), enamelin (ENAM), and ameloblastin (AMBN), which are all secreted by ameloblasts ^31^. Amelogenin exists in two isoforms, AMELX and AMELY, encoded on the non-recombinant regions of the X and Y chromosomes, respectively. By specifically detecting peptides derived from these isoforms using MS/MS, proteomics analysis allows for sex determination. Since human females have two X-chromosomes, they only express AMELX protein, whereas the male, having both an X- and a Y-chromosome, expresses both AMELX and AMELY proteins. Consequently, confident detection of AMELY-specific peptides confidently identifies male specimens, whereas the exclusive identification of AMELX-specific peptides does not preclude males and cannot conclusively point to female origin.

The typical workflow for MS-based sex identification currently involves sensitive and extensive liquid chromatographic (LC) separation coupled with high-resolution MS/MS analysis using a data-dependent acquisition (DDA) strategy. In DDA, the peptides are selected for fragmentation based on their relative MS signal intensities resulting in a bias towards sequencing of the most abundant peptide species ^32,33^. This acquisition scheme acquires MS/MS spectra throughout the entire LC gradient in a hypothesis-free manner, where the nature of the peptides doesn’t have to be known prior to data acquisition. This strategy, however, generates large amounts of data, from which the peptide precursors need to be identified using a database search engine, to match each experimental tandem mass spectra against in-silico generated spectra, based on a protein sequence reference database ^34^. The matching of MS/MS spectra to a peptide sequence is computationally time-consuming and requires proteomics skills and experience to analyse the outputs. This approach is suboptimal for the reproducible identification or quantification of low-abundant peptides, without a streamlined data analysis strategy. Alternatively, data-independent acquisition (DIA) has shown great potential for reproducible and sensitive quantitative analysis. In DIA, wide mass range windows, typically 25 Da, are selected for fragmentation. The windows successively cover the full mass range within a scan cycle, removing the stochasticity inherent to the DDA approach, and reducing the number of missing values. However, DIA tandem mass spectra are highly complex due to the co-fragmentation of multiple precursors, therefore requiring the use of an experimental library derived from previous analyses for the peptide identification from dental enamel. Attempts to standardise sex identification using a targeted MS data acquisition approach have been carried out ^35–37^. Parallel Reaction Monitoring (PRM) is a targeted proteomics technique based on high-resolution MS/MS instrumentation, typically with an Orbitrap or time-of-flight analyser ^38^. In PRM, a set of predefined precursor ions are specifically targeted in the mass spectrometer with high sensitivity and high selectivity. A PRM scan cycle consists of a full MS scan followed by quadrupole isolation and fragmentation of the targeted precursor ions in turn, whereby parallel monitoring of all fragments from each precursor ion individually can be achieved. This is a method of choice to perform reproducible quantitative analysis, where the subsequent data analysis mostly focuses on extracting and quantifying the ion signals without the need for a database search.

Here, we propose a standardised strategy that employs mass spectrometry for the assessment of the biological sex of human remains. We developed a fast, non-destructive, sensitive, robust, and quantitative MS-based assay for the determination of biological sex from archaeological dental enamel. Our workflow incorporates a micro-destructive acid etching enamel proteome extraction ^20^, followed by a proteolytic digestion-free ^29^ peptide purification strategy. We combined the fast extraction protocol with a fast LC-MS/MS acquisition method enabling the analysis of up to 200 samples per day (SPD) and presenting a semi-automated data analysis pipeline for reproducible and confident biological sex assignment. As the PRM targeted data acquisition approach allows for a highly reproducible quantitative MS analysis, we were able to model the monitored AMELX and AMELY peptide intensity relationships, predict AMELY peptide intensities when not detected, and obtain confident positive female identifications. This pipeline aims to simplify the entire analytical workflow, to ultimately increase the throughput and accessibility of palaeoproteomics-based sex identification studies for understanding population demographics, behaviours, and roles in past societies.

The method was first developed on 20 modern teeth and then tested on a set of 20 modern deciduous teeth, both of known sex. The developed methodology was initially applied to a set of 24 archaeological samples from Jordan, dating from ∼ 9000 BP, and showing poor morphological dental preservation, for assessing the applicability of the entire pipeline on ancient Holocene material. Finally, the scalability and robustness of the method were demonstrated through the analysis of > 300 teeth from the late Middle Ages, 1359-1360 CE, archaeological site of the Domplatz in Sankt Pölten, Austria.

## Results

### Selection of the target peptides for the development of a PRM assay

For the development of a targeted MS assay, it is crucial to identify the proteotypic peptides that can be consistently detected by MS. Yet, once enamel enters its maturation phase, the organic matter is degraded through proteolytic enzymes ^26^. This phenomenon in addition to the diagenetic degradation of the material and the harsh acidic peptide extraction leads to the recovery of unspecifically cleaved peptides. A cohort of eleven enamel samples from living individuals, 5 males and 6 females were analysed to investigate their enamel peptide composition. Enamel peptides were extracted using a 10% trifluoroacetic acid (TFA) etching, cleaned up on Evotips, and separated online on an Evosep One LC system using a 30 SPD gradient. The peptides were sequenced by higher-energy collisional dissociation (HCD) ^39^ with an Orbitrap Exploris 480 mass spectrometer operated in DDA mode to achieve high MS coverage of the extracted enamel peptides. The resulting raw MS/MS spectra were searched using MaxQuant against the human enamelome database (**Figure 1-A**).

**Figure 1:**
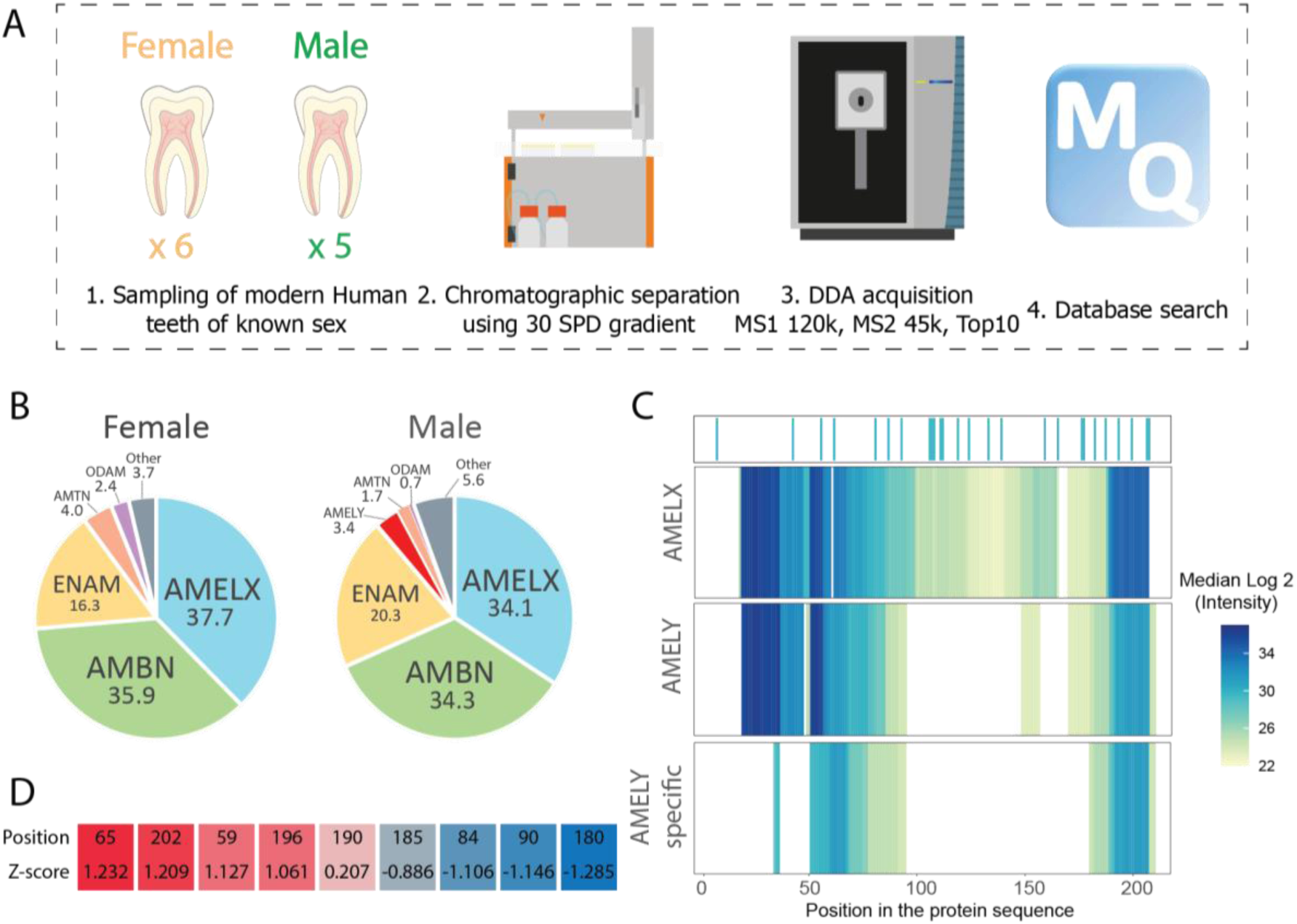
Dental enamel is composed of a few proteins, including amelogenin. A. Schematic representation of the workflow used for the selection of the target peptides for building the assay. B. Representation of the enamel composition for female and male individuals, calculated as PSM counts from the 11 modern individuals analysed in DDA. The numeric values represent the percentage of PSMs assigned to the given gene. C. Representation of the AMELX and AMELY coverage on each amino acid position calculated as the median log2 intensity of each position. Dark blue represents a site with high intensity while yellow shows sites with low intensity. The top plot shows the variant amino acid positions between AMELX and AMELY. The bottom plot (AMELY specific) displays the AMELY peptides based on protein grouping according to Occam’s razor principle. D. Table displaying the Z-score calculated for each amino acid position covered in both AMELX and AMELY using MS, to highlight the most intense sites. The Z-score is calculated as the subtraction of the site intensity of a given position with the average site intensity of all AMELY-specific positions, divided by their standard deviation.

This approach enabled a comprehensive overview of both the enamel proteome composition and its coverage using mass spectrometry (**Figure 1-B**). In all samples, the most abundant enamel proteins were AMELX and AMBN followed by ENAM (**Figure 1-B**). These proteins accounted for up to 90 % of the Peptide-Spectrum Matches (PSMs) and showed a good sequence coverage spanning from 90 % for AMELX, 68 % for AMBN to 20 % for ENAM. In total, 26 differential amino acid positions or sites exist between AMELX and AMELY in *Homo sapiens,* but only 9 of them were covered using tandem MS in both AMELX and AMELY with the proteolytic digestion-free approach. This partial coverage of both AMELX and AMELY could be due to the peptide length, chemical composition, or abundance, making them unfit for conventional LC-MS/MS analysis. To identify the most abundant peptides covering AMELY- and AMELX-specific sequence sites, the detected peptides were aligned and mapped to their gene positions and the PSMs were collapsed per charge state and modifications into stripped peptide sequences. The intensity for each amino acid position was then calculated as the sum of the intensities of all precursors containing that specific site (**Figure 1-C**).

Based on the AMELY coverage in the 5 male test samples, intensities across the measured amino acid positions were normalised by z-score (the average intensity of each site was subtracted by the average intensity and divided by the standard deviation of all measured sites) (**Figure 1-D**). Among these, four amino acid sites (AMELY-65, AMELY-202, AMELY-59, and AMELY-196) showed a high Z-score (>1) and were consequently further investigated for the differentiation between AMELX and AMELY. Various overlapping peptide sequences covering those sites of interest could be detected due to the non-specific proteolytic nature of the extracted peptides. To shortlist peptide candidates, filtering was applied to the peptide lists to select only peptide sequences with fewer than 12 amino acids and quantified across at least 80% of the samples. Moreover, peptides bearing modifications, except for methionine oxidation, were filtered out. After filtering, no peptides covering AMELY-202 or AMELY-196 remained.

Further manual filtering of the potential peptide targets using Skyline was carried out, selecting only precursors showing both MS1 and MS2 symmetric elution profiles as well as intense diagnostic fragment ions. In total five peptide precursors were selected for building the targeted assay: SIRPPYPSY (m/z = 540.2796 Th, z = 2), SIRPPYPSYG (m/z = 568.7904 Th, z = 2), SMIRPPY (m/z = 432.2258 Th, z = 2), SM(ox)IRPPY (m/z = 440.2233 Th, z = 2) and SM(ox)IRPPYS (m/z = 483.7393 Th, z = 2). Those targets were among the ones with the highest MS signal intensities covering the unique AMELX and AMELY sites of interest **(Figure S1)**.

### Development of a robust end-to-end strategy for sex identification

Here, we propose a robust pipeline for high-throughput analysis of dental enamel samples for confident sex identification (**Figure 2-A**).

**Figure 2:**
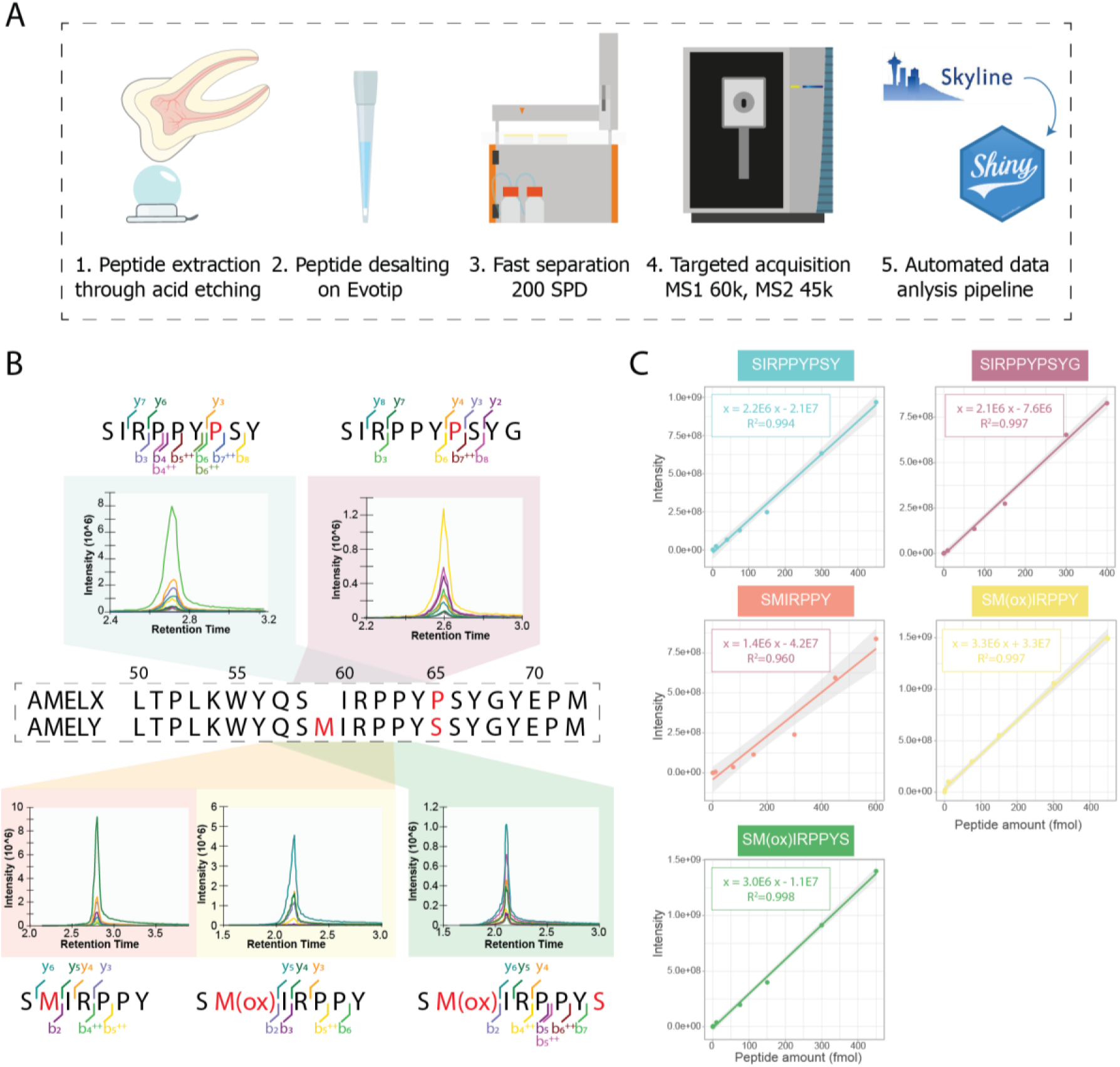
Five targets were selected for the building of the PRM assay. A. Schematic representation of the assaýs workflow. B. Display of the MS2 traces of the five selected target peptides. On the top, SIRPPYPSY and SIRPPYPSYG are specific to AMELX and cover both the amino acid positions 59 and 65. On the bottom, SMIRPPY, SM(ox)IRPPY, and SM(ox)IRPPYS are specific to AMELY, and all cover the amino acid position 59. SM(ox)IRPPYS also covers the position 65. The amino acids displayed in red are the variant amino acids between AMELX and AMELY. C. Linear regression of a dilution series of the target peptides ranging from 1 to 600 fmol for SMIRPPY and from 0.1 to 450 fmol for the other four target peptides. The grey areas represent the 95% confidence interval of the linear regression.

For peptide extraction, we relied on an acid etching strategy using 10% TFA, which minimises the alteration of the archeological remains and allows their morphological preservation ^20^. In this digestion-free approach, non-tryptic peptides are extracted along with the apatite mineral, the main constituent of the enamel matrix ^26^. The peptides are desalted and stored on Evotips that function as disposable pre-columns from which the peptides are automatically eluted online in the Evosep One LC system ^40^.

A dilution series of five specific peptides, ranging from (0.1 to 450 fmol) was analysed using a PRM method in the Orbitrap Exploris 480 and showed good chromatographic separation using the 200 SPD gradient (**Figure 2-B**). All of the selected targets demonstrated high linearity (**Figure 2-C**) over an intensity range covering three to five orders of magnitude. As a result, these peptide targets were considered suitable for quantitative analysis. Using a 4.5 minute effective gradient enables the analysis of 200 SPD without carryover across runs **(Figure S2)**.

The low number of peptide targets in the PRM acquisition method allowed for a high number of MS measurement points over the LC elution peak, guaranteeing precise quantification at the MS2 level, even though using a high-resolution method coupled with a short gradient allows for the analysis of 200 SPD. The quality of the measurement became apparent with the low coefficient of variation (< 7 %) across twelve technical replicates of pooled modern male samples (**Figure S3**) and demonstrates the possibility of performing precise quantification.

**Figure 3:**
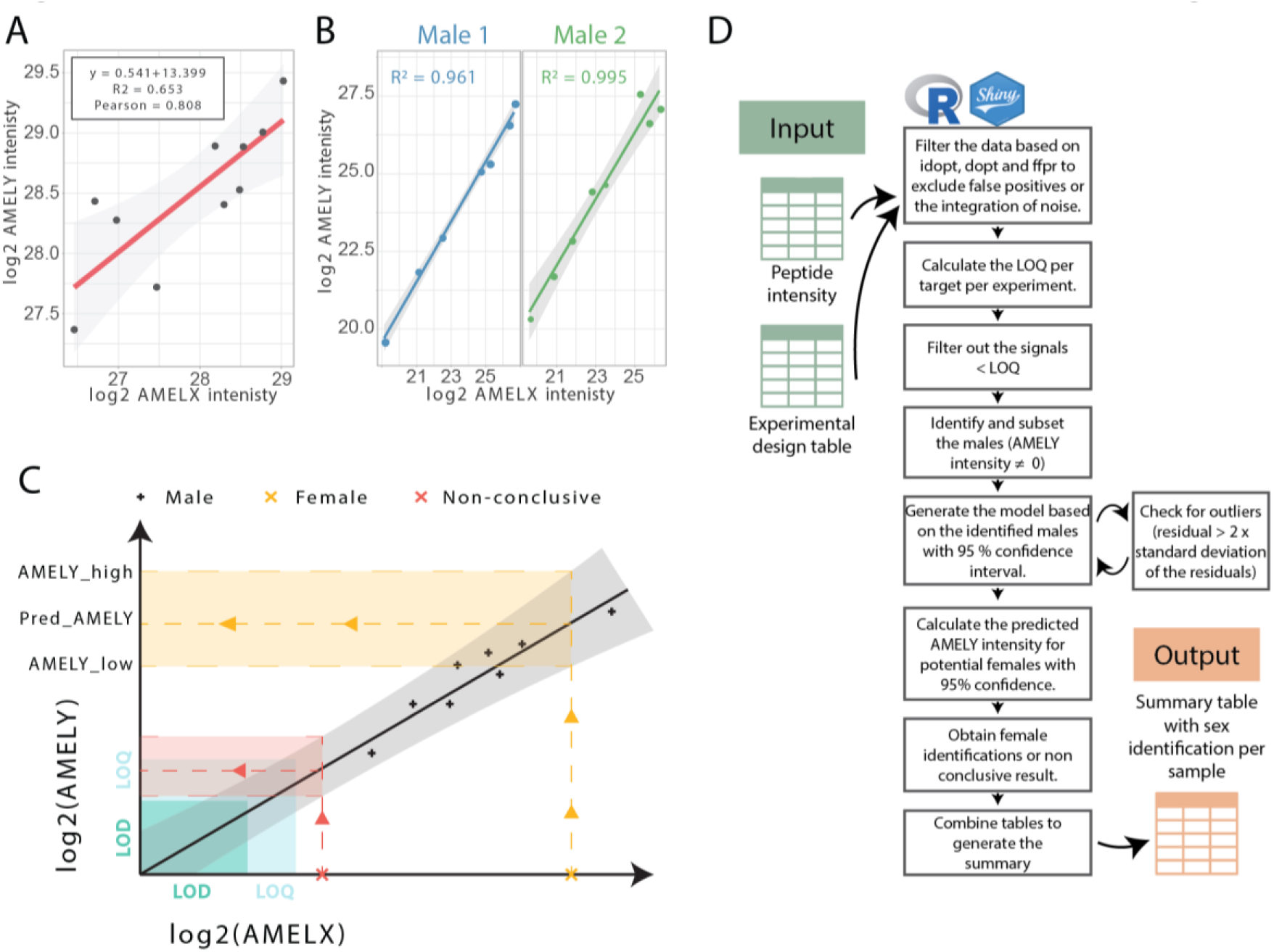
The linear correlation between AMELX and AMELY intensities is exploited for sex identification. A. Linear regression of the logarithmic transformed AMELY intensity as a function of the logarithmic transformed AMELX intensity in 10 modern male individuals. Each dot represents a different individual, the red line represents the linear model and the grey area represents a 95% confidence interval calculated using the standard error of the fitted values. B. Linear regression of the logarithmic transformed AMELY intensity as a function of the logarithmic transformed AMELX intensity for 2 different modern male individuals. Each plot represents a dilution series with each dot with a different input amount injected into the MS. The line represents the linear regression and the grey area represents a 95% confidence interval calculated using the standard error of the fitted values. C. Schematic representation of the sex identification process. The male individuals (black crosses) generate the linear model. Using the model, a predicted AMELY intensity is calculated from a measured AMELX intensity. If the output fits the imposed criteria, the sample is identified as a female (yellow cross), if not, the sample is given as non-conclusive (red cross). D. Flow chart of the data analysis pipeline.

Males expressing AMELY can be identified through the presence of AMELY targets. Yet, the absence of AMELY in our measurements cannot directly pinpoint those samples that are from females ^21^. Therefore, we aimed to develop a precise quantitative analysis pipeline to exploit the AMELX/AMELY peptide MS signal intensity relationships. That way, we can obtain confident female identification and prevent false-positive female identifications. To perform quantitative analysis, it is crucial to delineate the limit of detection (LOD) and most importantly, the limit of quantification (LOQ) of each peptide target. The limit of detection represents the lowest signal that can be detected in the mass spectrometer and the limit of quantification indicates the lowest MS signal that can be confidently quantified. To establish these, a dilution series of standard solutions containing the target peptides is run in the MS, prior to the analysis of the samples, using the same parameters as the samples. A linear regression analysis was performed on each target to determine the LOD and LOQ. Importantly, for obtaining accurate results, the standard dilution is specific to the MS set-up and time as it represents the performance of the instrument at a given time.

Efforts were made to automate the entire data analysis process, starting from the raw MS files to the final sex identification, where no prior proteomics experience is required. First, measured ion signals are extracted from the raw files using the freely available and open-source Skyline software ^41^. Secondly, using a graphical interface implemented in R-Shiny, the signals are further processed to obtain the identification of the biological sex of each sample. Before proceeding to the analysis, the extracted ion signals must be filtered to discard noisy signals or false identifications and ensure that only the signal specific to the targets is being extracted ^42^. This quality filter is based on the “Isotope dot product” > 0.9 (similarity between the measured isotopic abundance distribution and the theoretical one calculated based on peptide sequence), “Library dot product” > 0.9 (similarity between the measured transition peak areas and the reference ones from the spectral library) and “Peptide Peak Found Ratio” > 0.9 (coelution of the transitions with the precursor signal). Peptide targets in samples with intensities < LOQ are filtered out to ensure that all the measured targets are quantifiable. AMELX and AMELY intensities per sample are calculated as the sum of the intensities of their respective targets.

Using our male reference samples, we observed a linear correlation between AMELX and AMELY with an R-squared of 0.65 and a Pearson correlation of 0.808 (p-value = 0.0047) (**Figure 3-A**). As all quantified peptides were in the linear part of the intensity range (R2 > 0.96) (**Figure 2-C**) and an enamel dilution series also showed high linearity (R2 > 0.96) (**Figure 3-B**), we concluded that the variation observed using different individuals originates from inter-individual diversity. Using the modeled relationship, we were able to predict the AMELY intensity of samples where AMELY was not detected, and their respective confidence interval (calculated as 2x the standard error of the fitted values), based on the quantified AMELX intensity. The accuracy of the approach described above is highly dependent on the model used. Here we propose to let the user choose between using our pre-generated model based on AMELX and AMELY peptides from contemporary human samples or generating their model based on the males present in their sample cohort analysed. If the sample size allows, the second approach would provide a better model for the sex estimation. The experimentally based model can be refined in case of outliers. The user can set a threshold of the maximum number of values that are allowed to be removed if detected as outliers (residual (measured value - fitted value) is higher than 2x the standard deviation of all residuals). The algorithm will remove one outlier at a time and recalculate the model in between until either the maximum number of removable points is reached, or no outliers can be detected. That way, outliers will still be identified as males but not taken into account for the building of the model. With this pipeline, individuals are identified as males if signals corresponding to AMELY-specific peptides are detected above the LOQ. We would confirm a female attribution if the two following criteria are fulfilled: (i) the predicted AMELY intensity is above the AMELY LOQ and (ii) the lower limit of the predicted AMELY intensity is above the AMELY LOD (**Figure 3-C**). Using this approach, we can exclude the absence of detection of AMELY targets due to instrumental limitations.

### Use of a validation cohort for assessing the performance of the PRM-based assay

To develop the analytical assay presented in this project we selected a cohort of 20 modern human teeth of known biological sex (10 males and 10 females) from the Dentistry School of the Faculty of Health and Medical Sciences at the University of Copenhagen, and afterward validated it on a second cohort of 20 modern deciduous teeth from known sex, where 10 belonged to males and 10 belonged to females from the Ratón Pérez collection, hosted at the Centro Nacional de Investigación sobre la Evolución Humana in Burgos (Spain).

Reassuringly, 19 of 20 accounting for 95% of the samples from the validation cohort could accurately be assigned to their respective biological sex **(Figure 4-A)**. Sample 14, labeled as a male individual, was here identified as a female. Compared to all other test samples, sample 14 had the lowest AMELX intensity **(Figure 4-A)**. No trace of AMELY targets could be detected in the MS while the AMELX targets were present and showed a clean signal **(Figure 4-B)**, excluding the possibility of having filtered out AMELY targets that were not quantifiable (< LOQ). Additionally, we wanted to exclude the possibility of false identification of sample 14 due to (i) the absence of AMELY detectable peptides in sample 14 from the target list or (ii) a low MS signal, which would lead to the statistics used for the assay to be unfit.

**Figure 4:**
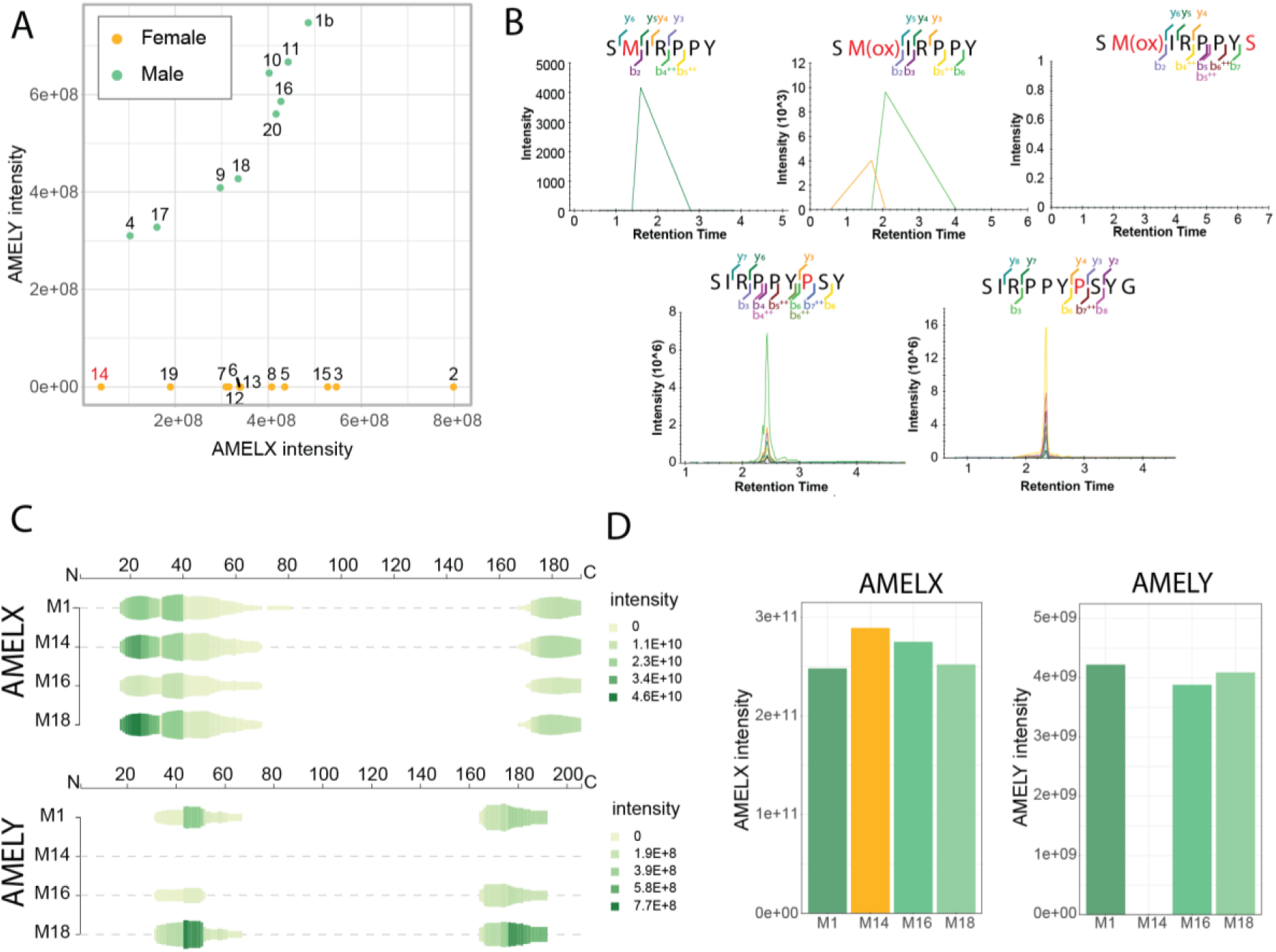
The assay was tested on a validation cohort. A. Overview of the sex identification of the 20 modern deciduous teeth from the Ratón Pérez collection represented as the raw AMELY intensity as a function of the raw AMELX intensity for each sample. Yellow dots are samples identified as females and green dots are samples identified as males. B. MS2 traces of each target peptide in sample 14 using the PRM acquisition strategy with a 200 SPD gradient. Top row: AMELY-specific targets. Bottom row: AMELX-specific targets. C. Overview of the AMELX (top) and AMELY (bottom) sequence coverage for samples 1, 14, 16, and 18 using peptigram ^47^ with DDA data acquired on a 30SPD gradient. D. Bar plot displaying the raw AMELX (left) and AMELY (right) intensity for each sample, with DDA data acquired on a 30SPD gradient. The protein intensities were calculated as the sum of the intensities of each precursor assigned to that protein.

Sample 14 was resampled and analysed using a DDA acquisition scheme on a 30SPD gradient, along with three other males (1, 16, and 18). AMELX could be identified in all the males and showed similar sequence coverage **(Figure 4-C)**. Yet, AMELY peptides could only be detected in samples 1, 16, and 18 and not in sample 14 **(Figure 4-C)**. This led to the conclusion that the targeted acquisition scheme was not responsible for the lack of detection of AMELY in sample 14. Moreover, the AMELX intensity from the DDA run of sample 14 was higher than any other male sample resampled **(Figure 4-D)**. We would then expect that, if AMELY were present in the sample, the mass spectrometer would be able to detect it. We can therefore conclude that the low AMELX intensity of sample 14 using the assay **(Figure 4-A)** is unlikely to cause the lack of detection of AMELY targets.

Altogether, we can only hypothesise that the false identification of sample 14 is due to either a miss labelling of the tooth or to the absence of AMELY from the sample itself, which could be caused by a deletion of the AMELY gene on the Y chromosome ^43–46^.

This test demonstrated the accuracy of the developed assay as well as the possibility of using it on deciduous teeth. Moreover, 95% of the samples were sexed accurately, showing that the method works on modern teeth. We also highlighted one of its main limitations: the inability to detect AMELY deletion, leading to a false female identification.

### Validation of the PRM-based assay on archeological samples and benchmark against traditional DDA-based strategy

Next, we applied the assay on archaeological samples using 24 dental remains from the Early Neolithic settlement Shkarat Mzaied (9.200-8.500 BP) in the Great Petra Region in Southern Jordan (**Figure 5-A**). The excavated skeletal elements showed poor preservation with bones in brittle condition and post-mortem damage on the dentition. With poor bone preservation and a high proportion of juveniles, osteological sex estimation is challenging, and out of 62 individuals from that site, only 16 could be assigned to a biological sex using osteological methods. 24 samples were selected for palaeoproteomics sex identification, consisting of 9 adults and 15 juveniles. Osteological-based sex identification was possible only for three of them.

**Figure 5:**
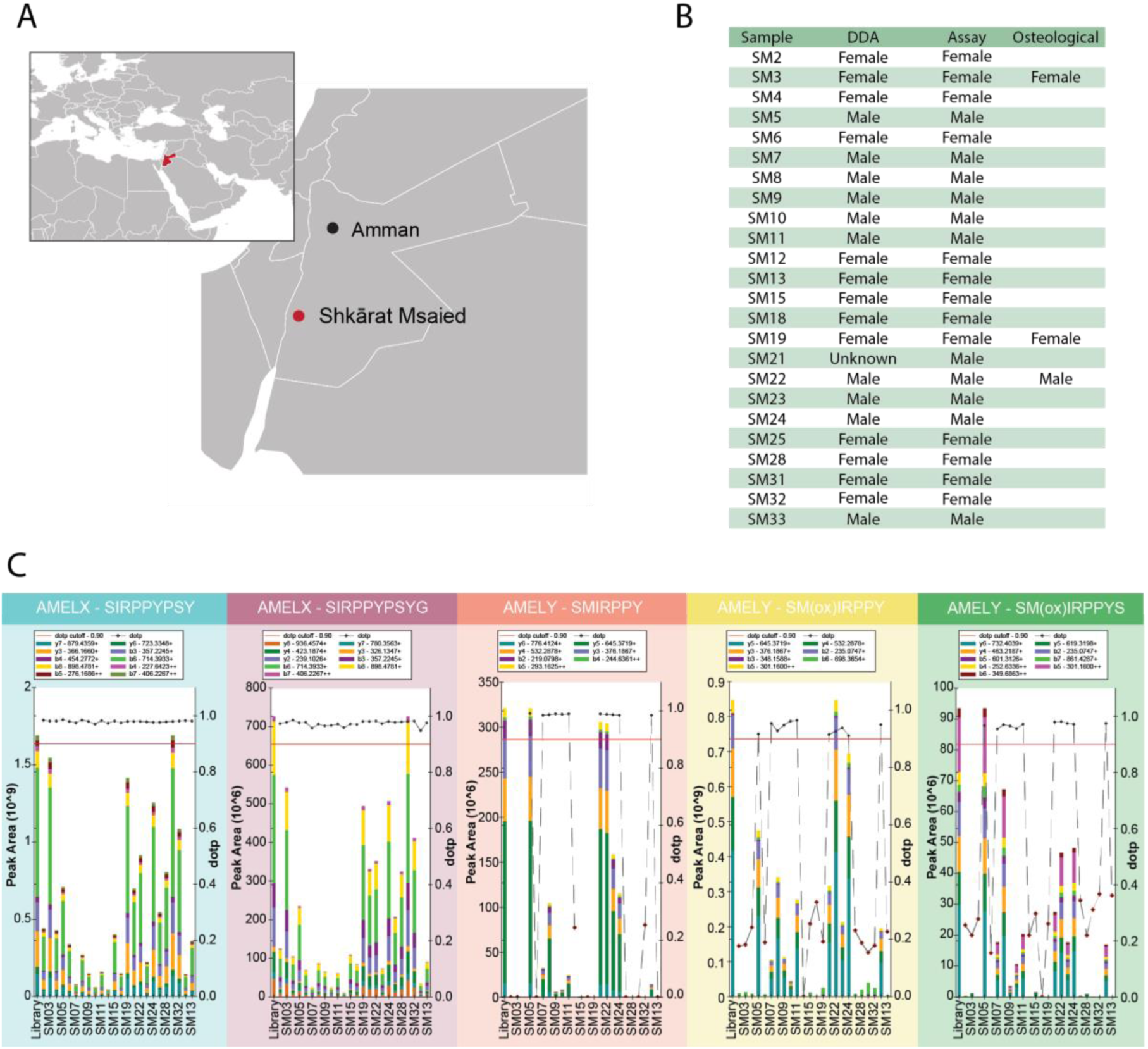
The PRM assay can identify the target peptides in archaeological samples. A. Location of the Shkarat Msaied excavation site on a map of Jordan. B. Table with the sex identification of the 24 archaeological teeth fragments from Jordan using a standard DDA approach, the developed PRM assay, and osteological estimations. C. Overview of the detection of the targets per sample. The bar plot represents the peak area and the various colours on the bars represent the proportion of each fragment ion detected. The correlation between this proportion with the one from the library (1st bar) is represented as black crosses (dopt). The red line represents the dopt cutoff, set at 0.9.

The teeth fragments were sampled and analysed twice, once using a previously published DDA-based method by Cappellini et al. ^29^, traditionally used for sex identification of ancient dental enamel specimens, and a second time, using the developed PRM assay strategy. The objective here was to control for the consistent detection of the selected targets in archaeological material. The DDA analyses were used as a means to validate the absence of any AMELY-specific peptides from the samples and not only the absence of the targets in the PRM approach.

Briefly, for DDA analysis, around 15 mg of enamel was demineralised at 4°C for 24h. The peptides were cleaned on StageTips, eluted into an MS plate, and separated online using a 77 minute nanoflow LC gradient. The raw LC-MS/MS files were searched using MaxQuant and the data analysis was performed through manual screening and validation of the identified sex-specific peptides. The biological sex was estimated by the absence or presence of AMELY peptides and ultimately led to the identification of 10 males and the tentative identification of 13 females. One individual remained unidentified due to the complete absence of enamel-specific proteins in its raw MS file, likely derived from a failed sampling.

Using the developed PRM assay, the biological sex of all 24 samples could be determined, leading to the identification of 13 females and 11 males **(Figure 5-B)**. The results obtained from the PRM assay were on par with the results using the previously published DDA-based method.

This experiment demonstrated that the peptide targets could be detected in all the samples **(Figure 5-C)**, enabling accurate sex identification of all of them. This also confirmed the choice of targets made, supporting their robustness and resistance through time periods. Altogether, the developed assay showed equivalent performance compared to the traditionally used sex identification methodology. At the same time though, the developed assay is far superior in terms of time efficiency in every experimental step: (1) sampling, (2) sample processing, (3) MS acquisition, and (4) data analysis. Altogether, using the assay, the results were obtained more than ten times faster and with less hands-on work.

### Demonstration of the high scalability of the assay with the analysis of > 300 archaeological samples

Performing fast and reliable sex identification on human teeth cohorts, including archaeological ones using a scalable workflow suited for high throughput applications, opens the possibility of standardising sex identification on larger sets of archaeological samples. As proof of concept, we utilised tooth samples from an extensive archeological excavation at the Domplatz in St. Pölten in Lower Austria (**Figure 6-A**). These samples were collected from the city cemetery ^48^. The sample set includes individuals from two mass graves: SG2 (N = 51) and SG 5 (N = 131), believed to have been used for burying victims of the bubonic plague pandemic, also known as the Black Death outbreak in 1359-60, along with 106 randomly selected individuals from various locations and time periods within the graveyard (mSG) (**Figure 6-B**). Additionally, 26 well-preserved skulls, intended for medical student training purposes, were retrieved from grave rearrangements and backfillings, contributing to the total sample size of 315 individuals. The total cohort was analysed in two different batches. Thus, two different standard lines were analysed. The specific LOQ threshold per target per batch was applied and the data were combined to generate the model used for sex assignment. Only one sample did not lead to a confident sex identification due to a low MS signal. Altogether, the 314 remaining samples led to the identification of 135 females and 179 males (**Figure 6-C**).

**Figure 6:**
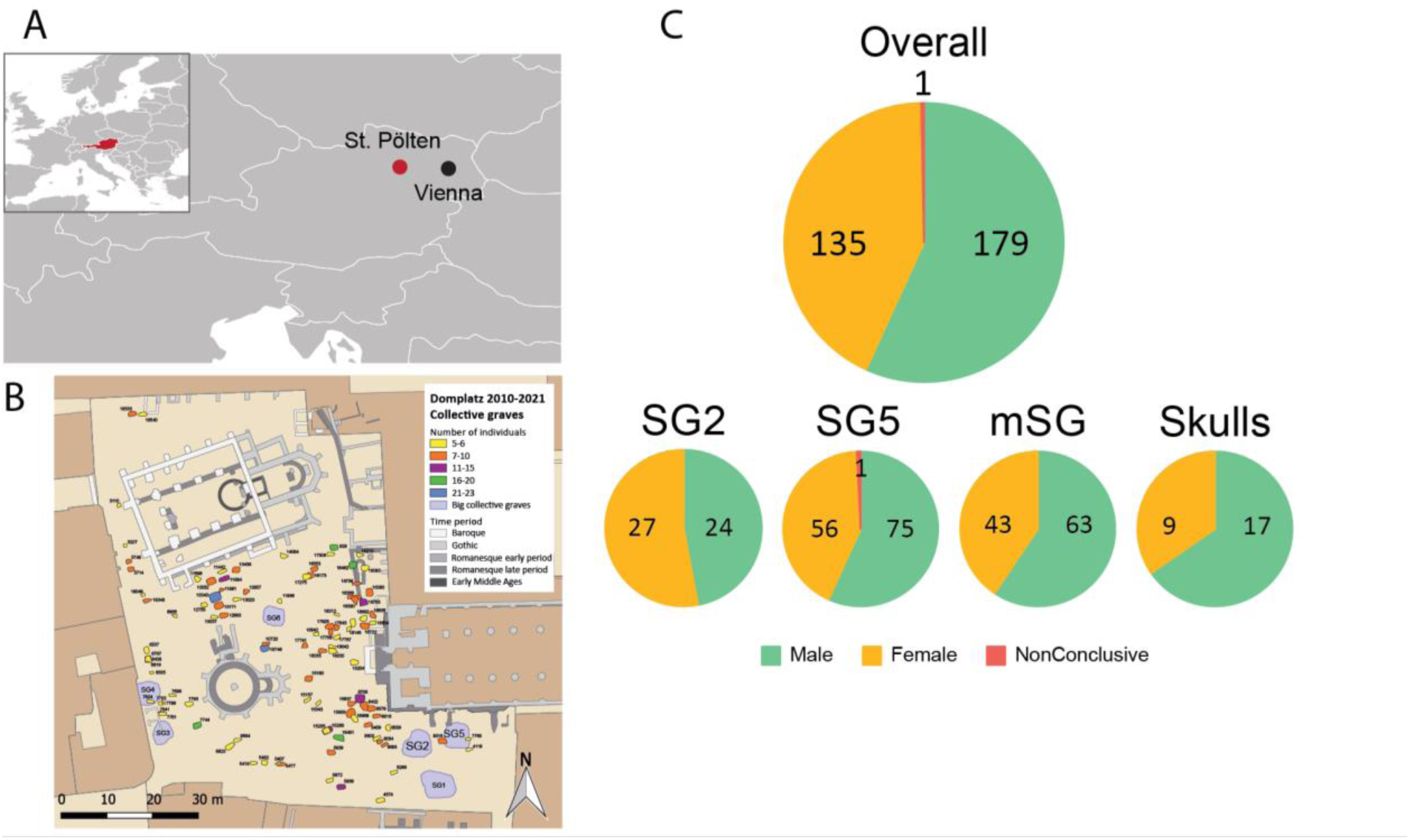
The biological sex of 315 individuals was assessed. A. Location of Sankt Pölten on a map of Austria. B. Map of the excavations on the Domplatz in Sankt Pölten, translated from ^48^ C: Pie charts showing the proportion of females, males, and non-conclusive identifications in the analysed cohort.

Obtaining the sex identification of the 315 samples required less than two days of MS time (1 day and 18 hours), including the analysis of the dilution series of the synthetic peptides with a total cycle time of 7.2 min, compared to almost 22 days (21 days 21 hours), with a cycle time of 100 min, which have been needed for the analysis using the DDA approach. Hence, with this fast chromatographic setup, accurate sex identification can be performed using one-tenth of the MS time previously required.

### Evaluation of the modelling strategy

In our assay, we use a linear model correlating the AMELX to AMELY intensity as a way to obtain confident female identification and remove the possibility of not detecting AMELY targets due to low abundance. The linear model can be built based on the data set studied. In that case, the cohort should be big enough to contain several male individuals so that a model can be built. Moreover, the more data points added for the construction of the model, the more likely those points will be spread across a wide intensity range, giving precision to the model. In case only a few samples are investigated, we also propose the option of using a predefined model built from modern individuals. Here, we wanted e to get a better understanding of the impact of the choice of the modelling strategy on the output of the assay.

We compared the different models generated through this study **(Figure 7**, **Figure S4)**. AMELX and AMELY peptides seemed to have different degradation rates. We observed that the “Dentist” and the “Ratón Pérez” models, both generated from modern teeth, were very similar (**Figure 7**). Conversely, the “St Pölten” and the “Jordan” models, both generated from archaeological material, showed different behaviour compared to the modern material. We observed a lower slope (0.5 for modern material, 0.7 for “St. Pölten” and 0.8 for “Jordan”) and a higher intercept (13.4 for modern material, 6.7 for “St. Pölten” and 4.8 for “Jordan”) in the modern material compared to the archaeological one. Additionally, the material from St. Pölten was more modern and showed less degradation than the one from Jordan. We thus observed an increase in the slope and a decrease in the intercept based on the age of the material.

**Figure 7:**
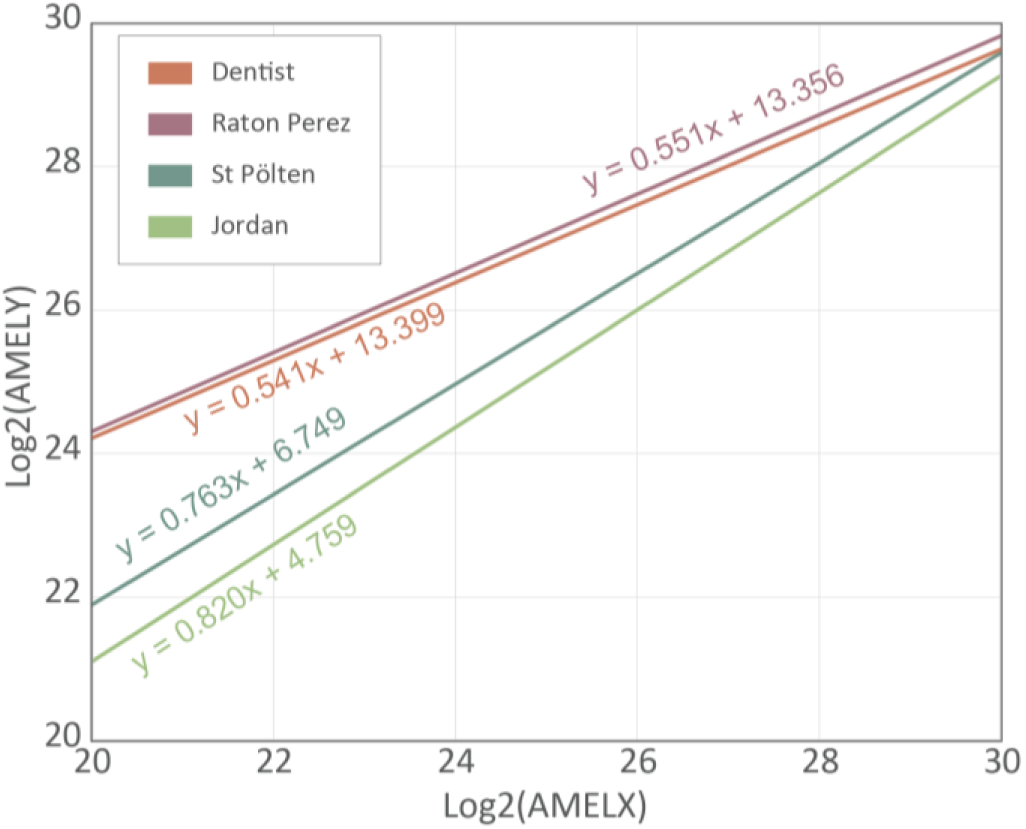
Dataset-specific models should be generated to avoid overestimation of the predicted AMELY intensities of potential females. Overlay of the four models generated during this study.

Applying a model based on modern material to archaeological samples would lead to an overestimation of the predicted AMELY intensities of potential females. This overestimation could then lead to false identifications, where samples would appear as females when they should appear as inconclusive. The effect also seems stronger at low intensity, where precision is needed to reach accurate sex identification. Altogether this evaluation highlights the need for generating data set-specific models.

## Discussion

Our research introduces a refined proteomics-based assay for the assessment of biological sex based on dental enamel. We provide an alternative to morphological-based or culturally-oriented sex inference by relying on biological data. The developed assay shows great potential for palaeoproteomics, where biological sex inference in archaeological contexts leads to a deeper understanding of past cultures. However, its adaptability renders it equally suitable for contemporary forensic investigations.

This PRM-based assay shows unprecedented scalability for molecular sex inference. To this date, no more than 75 archaeological individuals within one cohort have been studied using proteomics-based sex identification ^49^, with a throughput twice as high compared to Lugli et al ^50^, where 30 teeth were analysed (**Figure S5**). Currently, sex identification using proteomics is limited to a few key samples. Increasing the scale of the analysis is crucial to provide solid information for deepening our understanding of the sex-based power dynamic in societal structures, comprehensive studies on demographic patterns, burial practices, or even the prevalence of diseases between the sexes. Here, we take advantage of acid etching, a rapid and non-invasive peptide extraction method, in combination with a fast LC gradient and targeted MS acquisition method, and the addition of an automated data analysis pipeline to perform high throughput sex identification. With our assay, it is now possible to determine the sex of an individual from his dental enamel in less than half an hour. With the increased speed provided by the assay, we have demonstrated the feasibility of analysing > 300 samples in less than 2 days of MS time (**Figure S6**). The efforts on the standardisation of the entire pipeline, including the data analysis, can help democratise access to large-scale sex identification analyses using proteomics. The data analysis of > 300 samples took less than one day using the developed data analysis pipeline. This targeted MS assay’s ease of use allows its implementation in diverse laboratories, where an MS is available for data acquisition. Altogether, the high scalability, the reduced operational costs, and the minimal required expertise characterising this approach overcomes the current barriers limiting access to the study of sex identification through proteomics, ensuring wider accessibility to this information and applicability to large sample cohorts. Additionally, proteomics analysis of dental remains has so far been limited to high-profile samples, generating a bias in the areas of research. By democratising proteomics-based sex identification, we aim to remove that bias and allow the analysis of any cohorts of interest.

Additionally, improving experiment-specific workflows is a collaborative effort within the community, which enables more consistency and reproducibility across various analyses and laboratories. When established workflows undergo validation by the scientific community, it guarantees higher data reliability without additional user-driven optimisations, empowering researchers to make more effective use of the data from diverse studies. In the field of palaeoproteomics, standardised workflows would ascertain that differences observed among samples are genuinely sample-specific rather than from processing methods, which would allow for sample set wide comparison rather than reanalysing or resampling invaluable material. Some efforts have already been made to standardise palaeoproteomics workflows, with fully standardised MS-based protocols developed for species identification from bone fragments ^51^ and paint binder identification ^52^.

The developed statistical framework for positive female identification is pushing forward previous attempts. Here, we can identify both male and female individuals using a 95% confidence interval. Using a quantitative MS acquisition strategy, we were able to correlate the AMELX with the AMELY intensity to generate a model using male individuals, which enabled us to predict AMELY intensity when it was not detected. Using this framework, we can rule out that the absence of AMELY is due to the technique used (MS). However, our research has also demonstrated the limits of the assay, especially towards the deletion of the AMELY gene, leading to the absence of AMELY at the protein level. In that case, the assay will lead to false female identification necessitating confirmation via DNA analysis. Additionally, we are not able to predict how other biological conditions affecting the sex chromosome pair such as in trisomy for instance would affect the outcome of our assay.

Ultimately, the assay has been developed to fill a gap in palaeoproteomics, where a fast, accurate, and standardised approach was needed to perform large-scale biological sex inference. At the moment, the assay has only been tested on *Homo sapiens*, yet it could be possible to extend it to other species sharing the same target peptides for the differentiation of AMELX and AMELY. There are only a few publicly available amelogenin protein sequences on Uniprot. However, efforts have been made to translate primate genomes into protein sequences ^53,54^, and based on those translated sequences, we can confirm that the assay could be extended to the Catarrhini parvorder (old world monkeys) that encompasses the Cercopithecidae, the Hylobatidae and the Hominidae ^55^. Assessing the sex of other species would require the study of a population large enough to take advantage of the dynamic model built on the analysed dataset. An experimental model per species should be generated since there might be some divergence across species in the AMELX/AMELY intensity relationship. This limitation could affect the assay’s applicability to extinct species, especially hominins, where obtaining larger sample cohorts is challenging.

We have demonstrated that the targets could all be consistently detected in archaeological samples from challenging environmental conditions for protein preservation with the samples from Jordan. The selected targets showed outstanding time stability and could also be detected in much older data sets such as ∼800 ka *Homo antecessor* from Spain ^7^, 1.77 Ma Dmanisi *Homo erectus* ^7^, or 2 Ma South African *Paranthropus robustus* ^30^, paving the way to the analysis of much older specimens.

## Experimental procedures

### Origin of the samples

In total 20 modern human teeth (10 males, 10 females) were collected at the School of Dentistry, University of Copenhagen for the development of the assay. Additionally, 20 deciduous teeth (10 males, 10 females) were collected from the “Ratón Pérez” project ^56^ for the validation of the assay. The Ratón Pérez project has the approval from the Bioethics Commission of the University of Burgos.

24 teeth fragments from the Early Neolithic settlement Shkarat Mzaied (dated to the PPNB = 9.200-8.500P) in the Great Petra Region in Southern Jordan were collected for proteomics analysis.

315 archaeological samples originating from the excavations at the Domplatz in St. Pölten in Lower Austria ^48^ were collected from the city cemetery spanning the Medieval and Early Modern periods. The sample set includes Black Death victims estimated from 1359/1360 and collected from two mass graves: SG2 (N = 51) and SG 5 (N = 131), along with 106 randomly selected individuals from various locations and time periods within the graveyard. Additionally, 26 well-preserved skulls, intended for medical student training purposes were analysed.

### Sample preparation

#### ● Acid etching

The surface of the enamel was cleaned with MilliQ water and etched for a few seconds (∼2s) in 10 % TFA. The etching was discarded and the enamel surface was rinsed with MilliQ water before being dried on a piece of paper towel. The teeth were put in contact with 10 % TFA for 2 to 10 minutes. The sample was loaded on Evotip for subsequent LC-MS/MS analysis. When dirt/solid particles were observed, the etching solution was further clarified by filtration with a 0.45 μm filter plate (Millipore, Sigma–Aldrich) before Evotip loading to prevent blocking of the tip and the loss of the sample.

#### ● Enamel cutting

Enamel proteins were extracted based on the sample preparation protocol by Cappellini et al. 2019. A dental enamel fragment (∼ 15 mg) was separated from the dentine using a cutting disc mounted on a drill and demineralised in 500 µL of 10% TFA overnight at 4°C. This step was repeated two times in total until the enamel was completely demineralised. The peptides were desalted on C18 StageTips (Rappsilber, Mann, and Ishihama 2007), and eluted with 30 µL of 40% acetonitrile (ACN) 0.1% formic acid (FA) into a 96-well MS plate. The sample was concentrated and resuspended in 7 µL of 5% of ACN 0.1% TFA.

#### ● Synthetic peptides

The five peptide targets were synthesised and purchased from Genscript. Each 0.8 mg aliquot was resuspended in 1 mL of 0.1% formic acid. The standards were diluted to reach the following amounts per 20 μL of standard (Table 1):

**Table 1:** Amount of each target per standard in fmol.

| Standard | SMIRPPY (m/z 432.2258) | SM(ox)IRPPY (m/z 440.2233) | SM(ox)IRPPYS (m/z 483.7393) | SIRPPYPSY (m/z 540.2796) | SIRPPYPSYG (m/z 568.7904) |
| --- | --- | --- | --- | --- | --- |
| A | 10 | 2 | 10 | 10 | 10 |
| B | 7.5 | 1.5 | 7.5 | 7.5 | 7.5 |
| C | 5 | 1 | 5 | 5 | 5 |
| D | 2.5 | 0.5 | 2.5 | 2.5 | 2.5 |
| E | 1 | 0.2 | 1 | 1 | 1 |
| F | 0.1 | 0.02 | 0.1 | 0.1 | 0.1 |
| G | 0.01 | 0.002 | 0.01 | 0.01 | 0.01 |

20 μL of standard were loaded on Evotips following the manufacturer’s protocol. Each standard was run three times in the mass spectrometer prior to the analysis of the samples.

#### ● Estimation of the carryover

Potential carryover across runs was estimated by analysing a blank sample between every sample on the validation cohort.

### LC-MS/MS setup

#### ● PRM method

Peptides were online separated on an Evosep One LC system with the 200 SPD predefined gradient, using a commercial Evosep endurance column (4 cm x 150 μm x 1.9 μm, EV1107) packed with ReproSil-Pur C18 beads and connected to a stainless-steel emitter (30 μm, Evosep, EV1086) without temperature control. The Evosep One was coupled online through an electrospray source to an Orbitrap Exploris 480 (Thermo Fisher Scientific).

Spray voltage was set to 2.0 kV, heated capillary temperature at 275°C, and funnel RF at 40. The method consists of a full scan at 60,000 resolution at mass-to-charge (m/z) 200 in centroid mode from 400 to 600 m/z. A target mass filter was added based on the m/z ratio and charge state (z) of the target peptides. Five targets without intensity thresholds were added to the inclusion list (m/z 432.2258 - z = 2, m/z 440.223 - z = 2, m/z 483.7393 - z = 2, m/z 540.2796 - z = 2, m/z 568.7905 - z = 2). The precursor mass tolerance was set at 5 ppm. If a target is detected in the full scan, an MS2 scan is triggered with an isolation window of 0.7 m/z using 30% normalised collision energy. The fragment spectra resolution was set at 45,000 at m/z 200 with a maximum injection time of 86 ms and an automatic gain control (AGC) at 100%. Data was acquired in centroid mode.

#### ● DDA method with 30 SPD gradient

Peptides were separated Evosep One system using a commercial Evosep endurance column (15 cm x 150 μm x 1.9 μm, EV1106) packed with ReproSil-Pur C18 beads and connected to a stainless steel emitter (30 μm, Evosep, EV1086) without temperature control. The mass spectrometer was operating in positive mode with a spray voltage at 2.0 kV, heated capillary temperature at 275°C and funnel RF at 40. Full MS resolution was set at 60,000 at m/z 200 with a maximum injection time of 25 ms. The full scan mass range was set from 350 to 1400 m/z with an AGC target of 300%. The fragment spectra resolution was set at 45,000 with an injection time of 86 ms and a Top10 method. The AGC target value was set at 200%, the isolation window was set at 1.3 m/z and the collision energy was set at 30%. The data was acquired in profile mode.

#### ● DDA method with nanoflow chromatography

Peptides were separated on a liquid chromatographic system (EASY-nLC™ 1200) using an in-house packed column (15 cm x 75 μm, 1.9 μm) packed with C18 beads (Reprosil-AQ Pur, Dr. Maisch). The nanoLC was operated using a 77 min gradient. Buffer A was milliQ water, buffer B was 80% ACN, 0.1 % FA, and the flow rate was set at 250 nL/min. The mass spectrometer was operated in Data-Dependent Acquisition (DDA) in positive mode. Full MS resolution was set at 120,000 at m/z 200 with a maximum injection time of 25 ms. The HCD fragment spectra resolution was set at 60,000 with a maximum injection time of 118 ms and a Top10 method with 30 seconds dynamic exclusion. The isolation window was set at 1.2 m/z and the normalised collision energy (CE) at 30%. Injection blanks were run prior to every sample injection to limit the risk of carry over in the chromatographic column.

### Data analysis

#### ● PRM data analysis

Ions signals corresponding to the targeted peptide and fragment ions were extracted using Skyline (23.1.1.318) ^41^ with a custom-made template. Two custom made reports were exported: (i) Peptide intensities data (Peptide, Protein, Replicate, Peptide Peak Found Ratio, Normalised Area, Library Dot Product, Isotope Dot Product, Peptide Modified Sequence, File Name, Sample Type, Analyte Concentration and Concentration Multiplier) and (ii) Raw intensities data (Modified Sequence, Replicate, Precursor, Transition, Raw Times, Raw Intensities, Sample Type). The first report contains the measured peptide intensities to perform quantitative analysis and the second one contains the raw data from every extracted peak for the plotting of the MS traces with the R-Shiny app. A third report is generated containing the file name of each sample and linking each file to an experiment. A custom-made R-based script is used for performing the data analysis (**Figure 3-D**). The data is filtered based on the isotope dot product (≥ 0.9), the library dot product (≥ 0.9), and the peptide peak found ratio (≥ 0.9). The Standards were subset from the samples and the limit of detection and quantification were calculated based on linear regression. The limit of detection per target is calculated as 3.3 x (Residual Standard Error / Slope) and the limit of quantification is calculated as 10 x (Residual Standard Error / Slope). The targets with measured intensities below their respective limits of quantification are filtered out. AMELX and AMELY intensities are calculated as the sum of the intensities of their targets. Samples measured with a non-zero AMELY intensity are identified as males. The AMELY/AMELX intensity relationship is modelled by (1) generating a linear regression based on the prior male identifications from the dataset studied or by (2) using a predefined model based on the analysis of modern material generated in this study. For the first option, outliers can be controlled by allowing a maximum percentage of the data points to be removed. If a sample residual is > 2 * the standard error of the residuals in the model, that outlier is removed and the model is recalculated. This process is performed as long as there are either no outliers left in the dataset or the maximum number of points allowed has been removed. The confidence interval of the model is calculated with a 95% confidence as ± 2 * standard error of the fitted values. For each sample presenting a null AMELY intensity, the predicted AMELY intensity is calculated based on the model with the corresponding confidence intervals. AMELX and AMELY LOQ and LOD are calculated as the highest LOQ or LOD of the targets corresponding to AMELX or AMELY respectively. A sample is identified as female if the predicted AMELY intensity is above AMELY LOQ and if the lower limit of the predicted AMELY is above the AMELY LOD. If the sample does not fit one of these conditions, it will be annotated as “Nonconclusive” (**Figure 3-C**).

#### ● DDA data analysis

DDA files were searched using MaxQuant (v. 1.6.17.0 or 2.0.1.0) ^57,58^ against a custom-made database containing human enamel protein sequences (92 entries). Unspecific digestion was selected. No fixed modifications were set. Arginine to ornithine conversion, asparagine and glutamine deamidation, methionine and proline oxidation, serine and threonine phosphorylation as well as the conversion of glutamine and glutamic acid to pyroglutamate were set as variable modifications.

## Acknowledgements

C.K., R.S.P., P.P.M., M.M.T., E.C., and J.V.O. are supported by the European Union’s Horizon 2020 research and innovation programme under the Marie Sklodowska-Curie “PUSHH” training network, grant agreement No. 861389. Work at The Novo Nordisk Foundation Center for Protein Research (CPR) is funded in part by a donation from the Novo Nordisk Foundation (NNF14CC0001). R.P., P.P.M., G.B.T., J.V.O., and E.C. are supported by the European Research Council (ERC) under the European Union’s Horizon 2020 research and innovation programme (grant agreement No. 101021361). M.M.T. and M.M.P. receive funding from Project PID2021-122355NB-C33 financed by MCIN/ AEI/10.13039/501100011033/ FEDER, UE. M.M.T receives funding from The Leakey Foundation through Dub Crook support. M.M.P. has the support of the European Research Council within the European Union’s Horizon Europe (ERC-2021-ADG, Tied2Teeth, project number 101054659). The Ratón Pérez project has been financed by the Spanish Foundation for Science and Technology (FECYT) of the Ministry of Science, Innovation, and Universities and Fundación “la Caixa”, Caixabank. We would like to acknowledge Ingolf Thuesen and Moritz Kinzel from ToRs Institute at the University of Copenhagen, as project head and excavation director of the Shkarat Mzaied Neolithic Project, respectively.

## Data availability

The mass spectrometry proteomics data have been deposited to the ProteomeXchange Consortium (http://proteomecentral.proteomexchange.org) via the PRIDE partner repository ^59^ with the dataset identifier PXD049326.

Custom R-code for sequence assembly is available on GitHub at: https://github.com/ClaireKoenig/SexIdentification

## Author contribution

E.C. envisioned the study. C.K., A.M.V, E.C., and J.V.O. conceived the scientific strategy. C.K. designed and performed the proteomics experiments, analysed the data, and conceptualised the data analysis strategy. C.K., P.B., R.P., E.C., P.P.M., and G.B.T. contributed to the sampling of all the teeth used in this study. N.V.H. collected and provided the modern material from the dentistry school in Copenhagen. M.M.P. and M.M.T. provided the modern material for the “Ratón Pérez’’ collection. M.L.S.J. and S.M. provided and selected the teeth material from the Shkarat Mzaied excavation site. B.R., F.K., and C.G. collected and provided the samples from the Sankt Pölten excavation site. A.M.V. and C.K. conceived and developed the R-Shiny interface. C.K., A.M.V., and J.V.O. wrote the manuscript. All authors read, edited, and approved the final version of the manuscript.

## SUPPLEMENTARY MATERIAL

**Figure S1:**
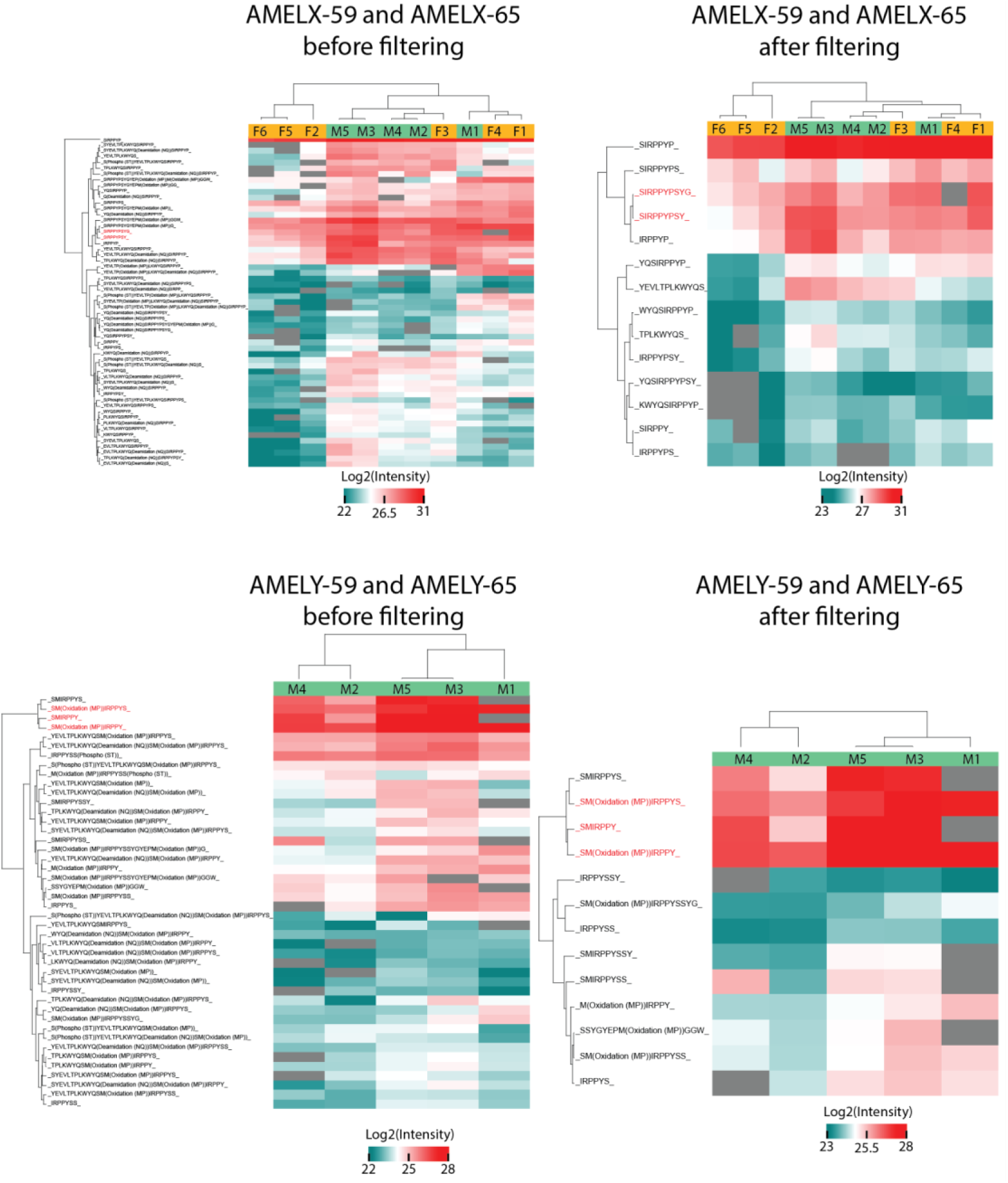
The target peptides are selected among the most abundant precursors. A. Hierarchical clustering of all the precursors covering the AMELX-59 and/or AMELX-65 amino acid positions in both male and female reference samples. Only precursors identified in at least 80% of the samples are displayed. B. Hierarchical clustering of all the precursors covering the AMELX-59 and/or AMELX-65 amino acid positions in both male and female reference samples after filtering. Precursors identified in less than 80% of the samples, with a sequence of more than 12 amino acids and containing modifications (except methionine oxidation) were filtered out. C. Hierarchical clustering of all the precursors covering the AMELY-59 and/or AMELY-65 amino acid positions in male reference samples. Only precursors identified in at least 80% of the samples are displayed. D. Hierarchical clustering of all the precursors covering the AMELY-59 and/or AMELY-65 amino acid positions in male reference samples after filtering. Precursors identified in less than 80% of the samples, with a sequence of more than 12 amino acids and containing modifications (except methionine oxidation) were filtered out. The precursors written in red are the target peptides that were selected for the PRM assay. The grey colour represents missing values.

**Figure S2:**
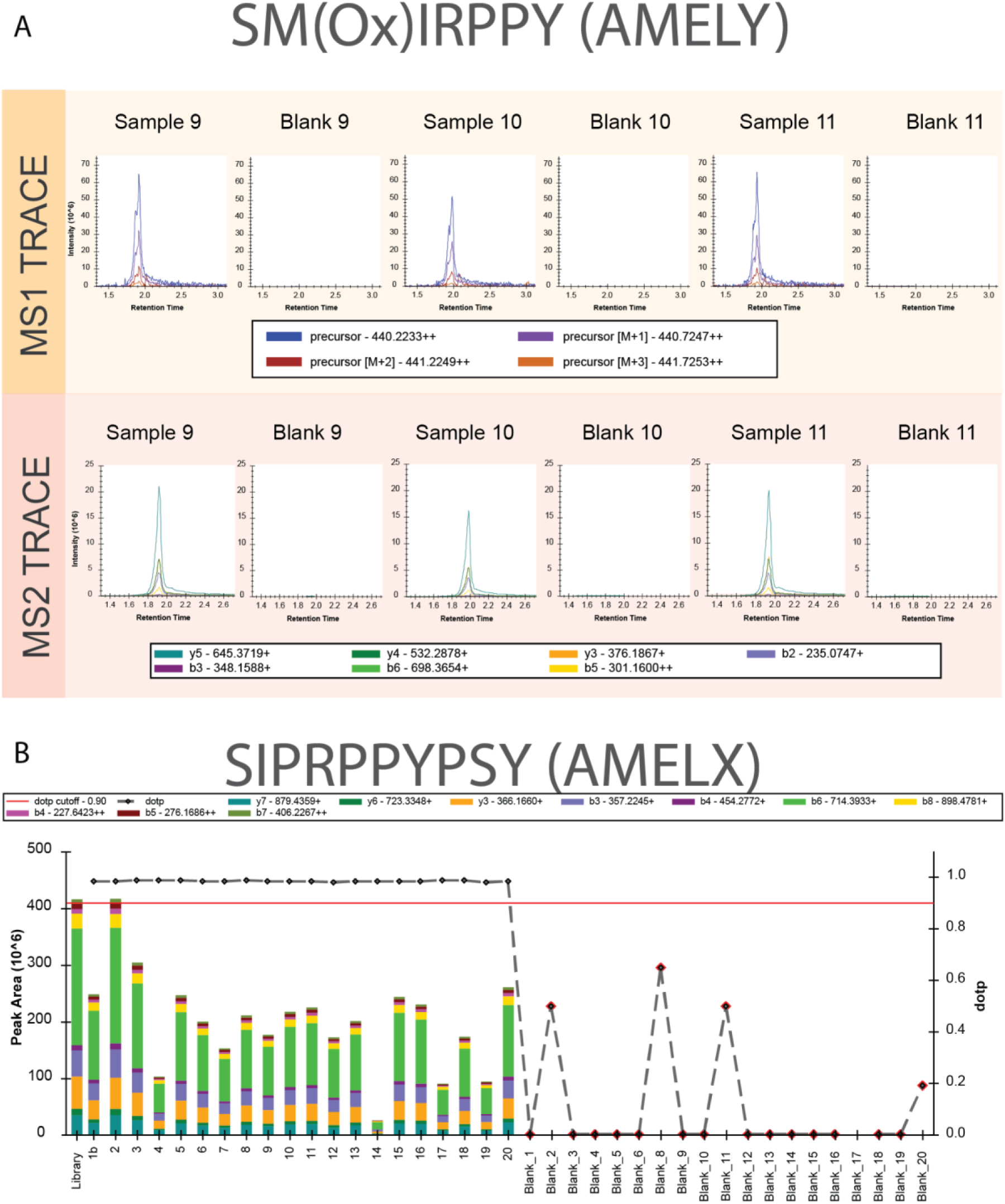
No carryover between samples could be observed. A. MS1 and MS2 traces of SM(ox)IRPPY in 3 male samples and 3 blanks run consecutively. The data originates from the validation cohort (Ratón Pérez), where blank samples were analysed in between samples. B. Bar Plot representing the peak area for SIRPPYPSY in the samples corresponding to the validation cohort (Ratón Pérez) and the blanks that were run between each sample. The black dots represent the dopt and the red line represents the dopt cutoff at 0.9.

**Figure S3:**
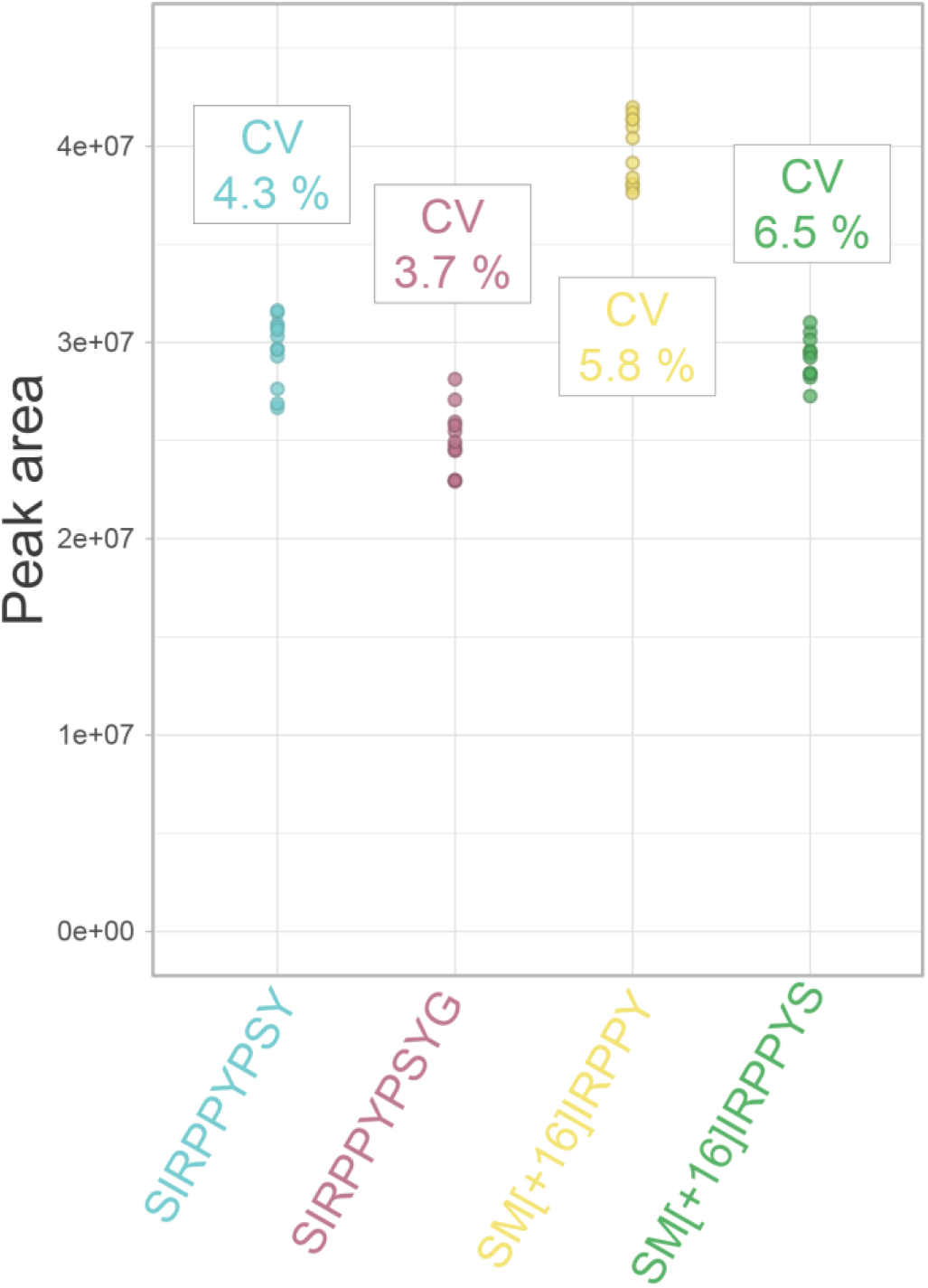
The PRM method shows good precision. Dot plot of the peak area (MS2) measured across 12 injection replicates of a pool of male enamel extract. The variation of SM(ox)IRPPY could not be measured since the target was absent from the samples.

**Figure S4:**
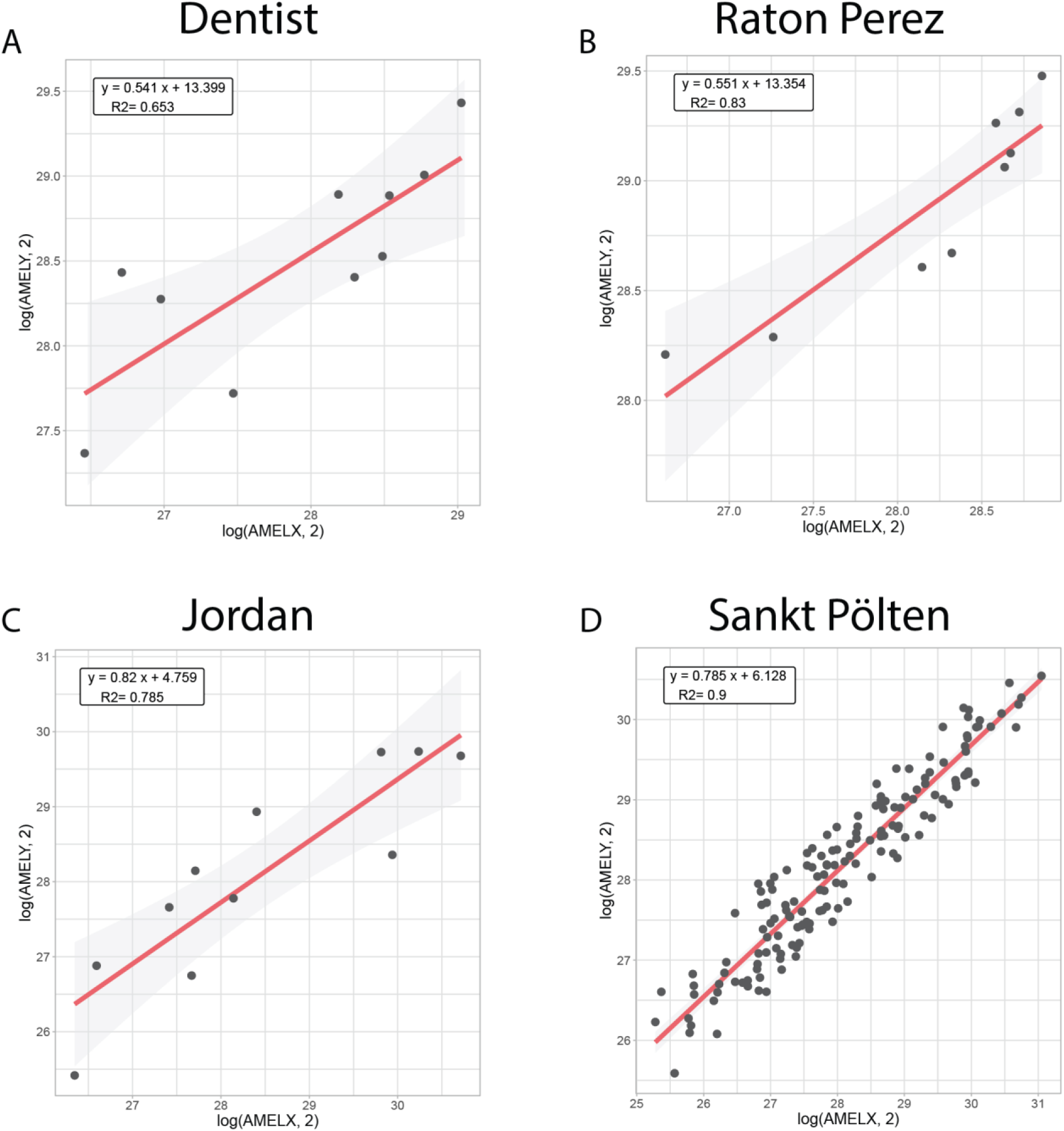
All studies cohorts showed linear behaviour between AMELY and AMELX intensities. A. Linear model generated from the samples from the dentistry school. B. Linear model generated from the validation cohort (Ratón Pérez samples). C. Linear model generated from the Jordan samples. D. Linear model generated from the samples from Sankt Pölten. *Figure S5:*#***Sex identification has so far mostly been limited to the analysis of a few high-profile samples.*** *The histogram represents the number of archaeological samples studied for sex identification using proteomics in the recent literature*.

**Figure S5:**
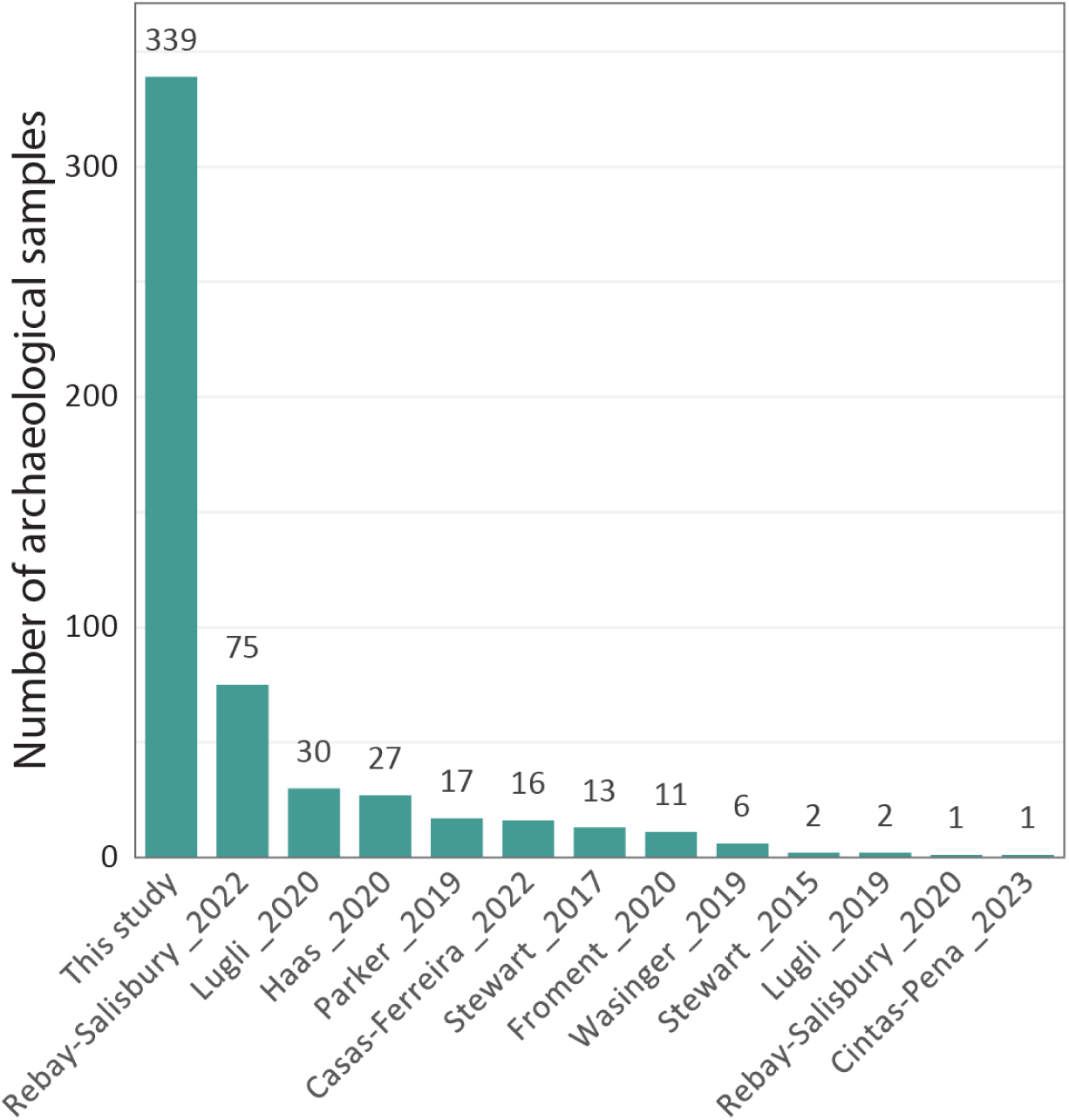
Sex identification has so far mostly been limited to the analysis of a few high-profile samples. The histogram represents the number of archaeological samples studied for sex identification using proteomics in the recent literature.

**Figure S6:**
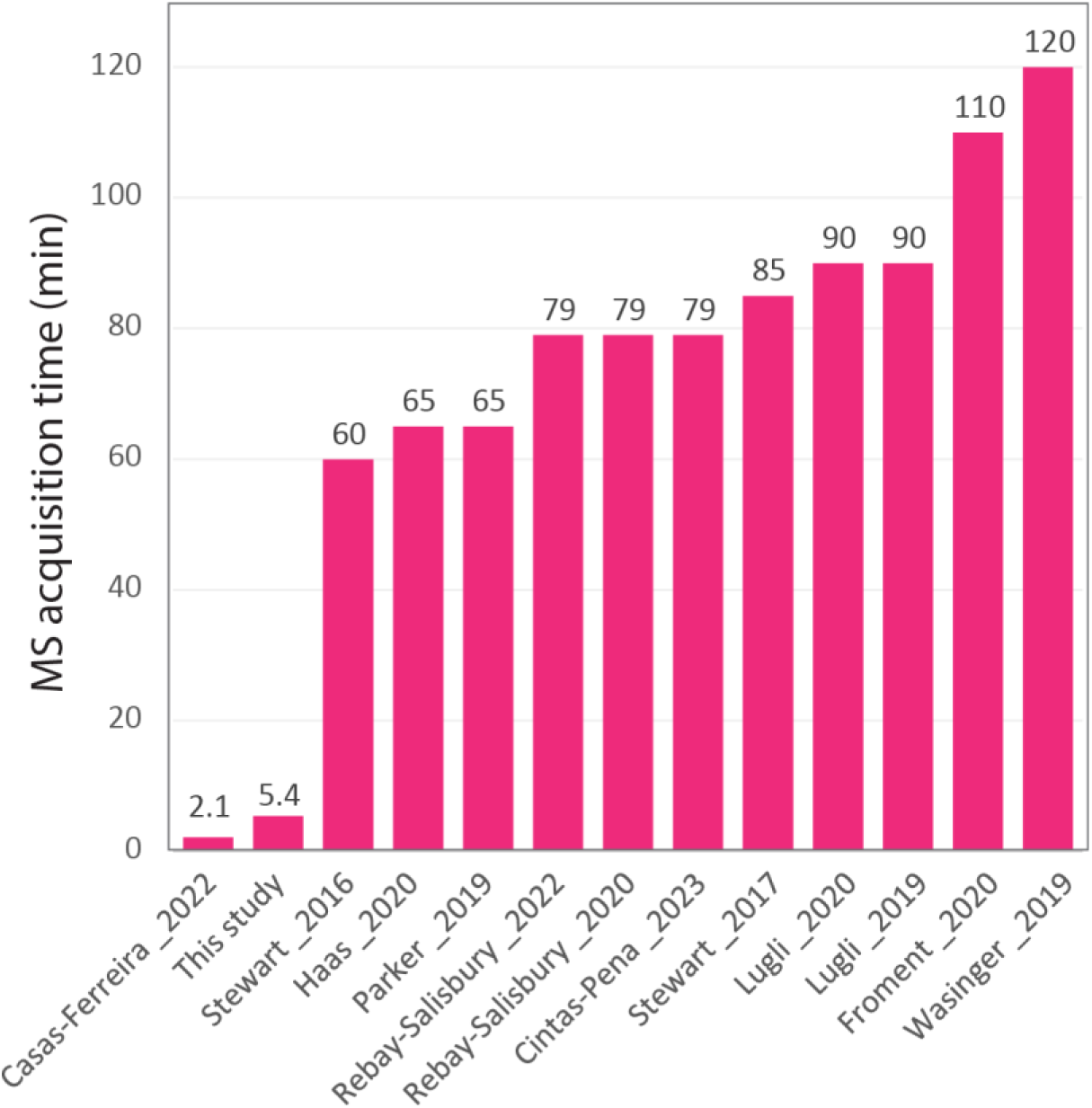
Sex identification has so far mostly been performed using extensive chromatographic separation. The histogram represents the length in minutes of the MS acquisition time used in the recent literature to perform sex identification. The MS acquisition time correlates with the length of the analysis and does not take into account the overheads of the LC system.

